# ADRA2A promotes the classical/progenitor subtype and reduces disease aggressiveness of pancreatic cancer

**DOI:** 10.1101/2024.03.12.584316

**Authors:** Paloma Moreno, Yuuki Ohara, Amanda J. Craig, Huaitian Liu, Shouhui Yang, Lin Zhang, Gatikrushna Panigrahi, Tiffany H. Dorsey, Helen Cawley, Azadeh Azizian, Jochen Gaedcke, Michael Ghadimi, Nader Hanna, S. Perwez Hussain

## Abstract

Pancreatic ductal adenocarcinoma (PDAC) manifests diverse molecular subtypes, including the classical/progenitor and basal-like/squamous subtypes, with the latter known for its aggressiveness. We employed integrative transcriptome and metabolome analyses to identify potential genes contributing to the molecular subtype differentiation and its metabolic features. Transcriptome analysis in PDAC patient cohorts revealed downregulation of adrenoceptor alpha 2A (ADRA2A) in the basal-like/squamous subtype, suggesting its potential role as a candidate suppressor of this subtype. Reduced ADRA2A expression was significantly associated with a high frequency of lymph node metastasis, higher pathological grade, advanced disease stage, and decreased survival among PDAC patients. *In vitro* experiments demonstrated that *ADRA2A* transgene expression and ADRA2A agonist inhibited PDAC cell invasion. Additionally, ADRA2A-high condition downregulated the basal-like/squamous gene expression signature, while upregulating the classical/progenitor gene expression signature in our PDAC patient cohort and PDAC cell lines. Metabolome analysis conducted on the PDAC cohort and cell lines revealed that elevated ADRA2A levels were associated with suppressed amino acid and carnitine/acylcarnitine metabolism, which are characteristic metabolic profiles of the classical/progenitor subtype. Collectively, our findings suggest that heightened ADRA2A expression induces transcriptome and metabolome characteristics indicative of classical/progenitor subtype with decreased disease aggressiveness in PDAC patients. These observations introduce ADRA2A as a candidate for diagnostic and therapeutic targeting in PDAC.

Graphical abstract

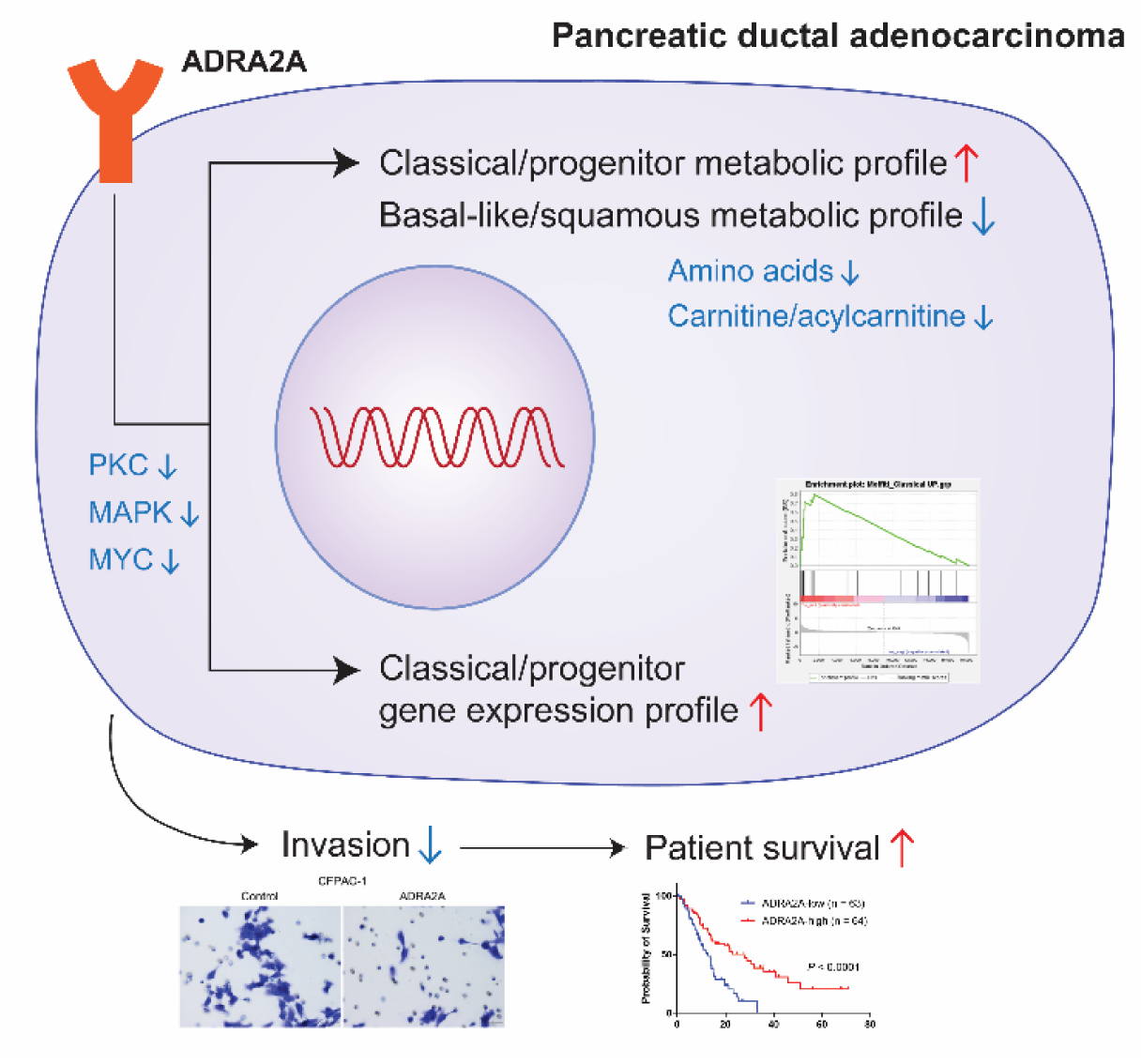

**Highlights:**

- ADRA2A is downregulated in the basal-like/squamous PDAC while its expression is maintained in the classical/progenitor PDAC subtype
- Upregulated ADRA2A expression correlates with improved PDAC survival and reduced invasion in PDAC cells
- Upregulated ADRA2A downregulates the MYC signaling pathway and promotes the classical/progenitor gene expression profile
- Upregulated ADRA2A induces a unique metabolic signature characterized by diminished amino acid and carnitine/acylcarnitine metabolism, resembling the classical/progenitor PDAC subtype

## Introduction

Pancreatic cancer continues to be one of the most lethal malignancies, boasting a dismal 5-year survival rate of merely 12% (1). Pancreatic ductal adenocarcinoma (PDAC) encompasses over 95% of all pancreatic malignancies, reigning as its most prevalent form (2). Within the landscape of PDAC research, distinct molecular subtypes have surfaced, each characterized by unique biological attributes holding implications for patient survival (3–5). *Bailey* et al. identified four molecular subtypes: aberrantly differentiated endocrine exocrine (ADEX), pancreatic progenitor, squamous, and immunogenic (3). The ADEX and pancreatic progenitor subtypes exhibit upregulated transcriptional networks containing various transcription factors associated with pancreatic development and the differentiation of exocrine and neuroendocrine lineages (3). The squamous subtype emerges as an aggressive form of PDAC, distinguished by specific traits such as metabolic reprogramming, proliferation, inflammation, hypoxia, squamous differentiation, and MYC activation (3). Recently, researchers divided PDAC into two major subtypes: ‘classical/progenitor’ and ‘basal-like/squamous’ subtypes (6). The ‘classical/progenitor’ subtype is more closely related to normal pancreatic tissue, reflecting its endodermal-pancreatic identity. In contrast, the ‘basal-like/squamous’ subtype shows characteristics associated with squamous differentiation (7).

Metabolic reprogramming, a key characteristic of cancer (8–10), plays a significant role in PDAC (10–12). Metabolic subtyping has revealed diverse metabolic profiles among different molecular subtypes of PDAC. The classical/progenitor subtype potentially favors lipogenesis, while the basal-like/squamous subtype shows a preference for glycolysis and amino acid metabolism (13–17). However, the intricate relationship between metabolic subtype and molecular subtype in PDAC remains inadequately explored due to limitations such as small sample sizes. Building upon our prior identification of SERPINB3 as an oncogenic driver of the basal-like/squamous subtype (17), this study aims to investigate the association between gene expression and metabolic adaptations in the development of the classical/progenitor subtype in PDAC. Our findings shed light on the role of that ADRA2A upregulation plays in shaping the metabolic landscape of the classical/progenitor subtype in PDAC.

## Materials and Methods

### Ethics Statement

Pancreatic tissues were acquired from surgically resected PDAC patients at the University of Maryland Medical System (UMMS) in Baltimore, MD, under the purview of an NCI-UMD resource contract, and from the University Medical Center Göttingen (Göttingen, Germany). Board-certified pathologists assessed PDAC histopathology. The use of these clinical specimens in our research in our study underwent review by the NCI-Office of the Human Subject Research Protection (OHSRP) at the NIH (Bethesda, MD) (Exempt#4678). All participants provided informed written consent, and all procedures adhered strictly to ethical standards and the principles outlined in the 1975 Declaration of Helsinki, as revised in 2008.

### PDAC cohorts

We utilized gene expression datasets from the “Bailey” cohort (GSE36924) (3), the “Moffitt” cohort (GSE71729) (4), and the NCI-UMD-German cohort (GSE183795) (18) to identify genes associated with PDAC molecular subtypes. Comparative analysis between basal-like/squamous and classical/progenitor subtypes was conducted using Partek Genomics Suite 7.0 (Partek Inc., Chesterfield, MO). For the integrated transcriptome and metabolome analyses, we used RNA sequencing data from PDAC tumors in the NCI-UMD-German cohort (GSE224564) (17).

### Cell lines and culture condition

Human PDAC cell lines were obtained from American Type Culture Collection (ATCC, Rockville, Maryland). Authentication of all cell lines through short tandem repeat (STR) profiling was performed within the past three years at ATCC. All experiments were conducted using mycoplasma-free cells. For CFPAC-1 (RRID:CVCL_1119) and Capan-1 (RRID:CVCL_0237) cell lines were cultured in IMDM supplemented with 10% FBS and 1% penicillin-streptomycin. The Capan-2 (RRID:CVCL_0026) cell line was cultured in McCoy’s 5A (Modified) with the same supplements. The remaining PDAC cell lines (AsPC-1; RRID:CVCL_0152, BxPC-3; RRID:CVCL_0186, MIA PaCa-2; RRID:CVCL_0428, PANC-1; RRID:CVCL_0480, Panc 10.05; RRID:CVCL_1639, SU.86.86; RRID:CVCL_3881) were cultured in RPMI 1640 with GlutaMax^TM^, 10% FBS, and 1% penicillin-streptomycin. All cell cultures were maintained in a humidified incubator with 5% CO2 at 37°C. All cell culture reagents were purchased from Thermo Fisher Scientific (Waltham, MA).

### ADRA2A overexpression after lentiviral infection

To establish stable cell lines overexpressing *ADRA2A*, the *ADRA2A* construct (EX-Z5688-Lv103) and the corresponding empty vector control (EX-NEG-Lv103), both sourced from Genecopoeia (Rockville, MD), were employed. Lentiviral particles were generated by transfecting 293T cells with the lentiviral expression vectors and the Lenti-Pac^TM^ HIV Expression Packaging system, also from Genecopoeia. To obtain stable clones of PDAC cells (Panc 10.05 and CFPAC-1), selection was carried out using 4 μg/ml puromycin (Thermo Fisher Scientific).

### RNA sequencing

Quadruplicate RNA sequencing was performed using total RNA extracted from the human PDAC cell line (CFPAC-1 +/− ADRA2A). The PDAC cells were cultured in IMDM supplemented with 10% FBS and 1% penicillin–streptomycin for 72 hours before RNA extraction. Libraries were constructed by the Sequencing Facility at NCI-Leidos using the TruSeq Stranded mRNA Kit (Illumina, San Diego, CA) and sequenced in a paired-end manner on NextSeq (Illumina) with 2 x 101 bp read lengths. This generated approximately 32 to 60 million paired-end reads, with a base call quality of ≥ Q30. The fastq-format sequence reads were aligned to the human reference genome hg38 using STAR and RSEM to obtain gene expression data, reported as transcripts per million with FPKM mapped reads. We performed differential expression analysis using DESeq2. Enrichment analysis, covering established pathways and datasets, was conducted using Ingenuity pathway analysis (IPA, QIAGEN, Venlo, Netherlands) and Gene Set Enrichment Analysis (GSEA). The RNA sequencing data have been deposited in the NCBI’s Gene Expression Omnibus (GEO) database under accession number GSE259390.

### Quantitative real-time PCR for *ADRA2A*

We used the High-Capacity cDNA Reverse Transcription Kit (Thermo Fisher Scientific) to synthesize first-strand cDNA from total RNA. Quantitative real-time PCR (qPCR) assays were conducted using Taqman probes (Thermo Fisher Scientific) for *ADRA2A* (Hs00265081_s1) and *GAPDH* (Hs99999905_m1).

### Metabolic profiling and data analysis of PDAC

Tumor sample metabolic profiling was conducted by Metabolon Inc. (Morrisville, NC) following their standard protocol (19–22). Metabolon’s untargeted metabolic platform employs two separate ultra-high performance liquid chromatography/tandem mass spectrometry (UHPLC/MS/MS) injections and one gas chromatography/mass spectrometry injection for each sample, enabling the measurement of all metabolites. The resulting dataset from the NCI-UMD-German cohort (n = 50) was normalized across samples before delivery (23,24), following Metabolon’s established protocol. The normalized relative abundance levels for each metabolite were collated and utilized for subsequent data analysis. The dataset was deposited previously.

The same standardized protocol was applied to conduct metabolic profiling of the human PDAC cells. Quadruplicate metabolic profiling of the human PDAC cells was conducted. The PDAC cells (Panc 10.05 +/− ADRA2A; 2 x 10^6^ cells) were seeded on a 100 mm dish and were cultivated in RPMI 1640 medium with GlutaMax^TM^ supplemented with 10% FBS and 1% penicillin–streptomycin for 72 hours. Subsequently, cell pellets were collected, preserved at –80 °C, and sent to Metabolon Inc. For enrichment analysis, we used MetaboAnalyst 5.0 (https://www.metaboanalyst.ca) (25).

### Metabolome analysis of PDAC cells with a MYC inhibitor

The absolute concentration of 116 metabolites was analyzed in various cell lines using the metabolome analysis package “Carcinoscope” provided by Human Metabolome Technologies, Inc. (HMT) (Boston, MA). For metabolite extraction, cell extracts were obtained following the standard manufacturer’s protocol (26). PDAC cells (BxPC-3; 6 x 10^5^ cells, Panc 10.05; 6 x 10^5^ cells) were seeded on a 60 mm dish and incubated in RPMI 1640 medium with GlutaMax^TM^ supplemented with 10% FBS and 1% penicillin–streptomycin, with or without a MYC inhibitor (100 μM, 10058-F4) for 72 hours in an incubator. We purchased 10058-F4 from MedChemExpress LLC (Monmouth Junction, NJ, HY-12702), and its corresponding solvent, dimethyl sulfoxide (DMSO), was used as the control. After removing the medium from the dish, cells were washed with 5% mannitol solution. Then, 400 μl of methanol was added and incubated at room temperature for 30 seconds. Subsequently, 275 μl of Internal Standard Solution was added and incubated at room temperature for another 30 seconds. One milliliter of the extracted solution was transferred to a 1.5 ml microtube and centrifuged at 4°C for 5 minutes (2300 x g). Then, 350 μl of the supernatant was transferred into a centrifugal filter unit and centrifuged at 4°C for 5 hours (9100 x g). Filtered samples were stored at –80°C until shipping. Metabolome analysis was performed by HMT using capillary electrophoresis mass spectrometry (CE-MS). IPA and MetaboAnalyst 5.0 (https://www.metaboanalyst.ca) (25) were used for the enrichment analysis.

### Invasion assay

24-well Falcon^®^ Cell Culture Insert (Corning, Glendale, AZ) and Matrigel Basement Membrane Matrix (#354234, Corning) were used for invasion assay. The upper chamber membrane was coated with 100 μl of Matrigel matrix coating solution (Matrigel matrix: coating buffer = 1:39) for 2 hours, following the manufacturer’s guidelines. The lower chamber was filled with 750 μl of medium containing 10% FBS, while the upper chamber was loaded with PDAC cells (Panc 10.05; 10×10^4^ cells, CFPAC-1 5×10^4^ cells) in 500 μl of serum-free medium. The cells were then incubated for 48 hours in a humidified incubator at 5% CO2 and 37°C. Following incubation, invaded cells were fixed using 100% methanol (Thermo Fisher Scientific) and subsequently stained with crystal violet solution (MilliporeSigma, Burlington, MA). Finally, the cells were counted. To examine the effect of clonidine (MilliporeSigma, C7897), 5 nM of the compound (final concentration) was added into the lower chamber.

### Immunofluorescence

PDAC cells were seeded on 8-well Nunc^TM^ Lab-Tek^TM^ II CC2^TM^ Chamber Slides (Thermo Fisher Scientific). Following PBS washing, cells were fixed in 4% paraformaldehyde in PBS at room temperature for 10 minutes, then rinsed twice and permeabilized with 0.2% Triton X-100 in PBS for 20 minutes. After another round of PBS rinsing, cells were blocked in a solution containing 2%BSA (MilliporeSigma) and 10% normal donkey serum (Thermo Fisher Scientific) for 1 hours at room temperature. Subsequently, cells were incubated overnight with rabbit polyclonal anti-ADRA2A antibody (Thermo Fisher Scientific, Catalog #14266-1-AP, 1:300). Following three washes with PBS for 15 minutes, cells were incubated with donkey anti-rabbit IgG (H+L) highly cross-adsorbed secondary antibody, Alexa Fluor^TM^ plus 594 (Thermo Fisher Scientific, A32754, 1:500) for 1 hour in the dark and at room temperature. After 15-minute wash, cells were stained with DAPI (Thermo Fisher Scientific, 62248, 1:1000) for 30 minutes, and mounted using ProLong™ Glass Antifade Mountant (Thermo Fisher Scientific). All samples were photographed under the same conditions using Zeiss LSM780 (ZEISS, Oberkochen, Germany).

### Immunohistochemistry

Four μm thick paraffin-embedded tumor sections, along with adjacent nontumor sections, were subjected to overnight incubation at 4°C with a rabbit monoclonal anti-ADRA2A antibody (Thermo Fisher Scientific, Catalog #14266-1-AP, 1:200). Signal amplification was facilitated using the Dako envision + system-HRP labeled polymer anti-rabbit antibody (Agilent Technologies, Santa Clara, CA). Color development was achieved using 3,3’-diaminobenzidine (DAB, Agilent Technologies). Immunohistochemistry (IHC) was evaluated by assigning intensity and prevalence scores (17,22). Intensity scores ranged from 0 to 3, representing negative, weak, moderate, or strong expression levels. Prevalence scores ranged from 0 to 4, indicating the percentage of cells showing ADRA2A expression (<10%, 10–30%, >30–50%, >50–80%, and >80%). The overall IHC score was calculated by multiplying the intensity and prevalence scores.

### Immunoblotting

We lysed PDAC cells by RIPA Lysis and Extraction Buffer from Thermo Fisher Scientific, supplemented with cOmplete™ Protease Inhibitor Cocktail (MilliporeSigma, 4693116001). Protein extracts underwent electrophoresis under reducing conditions on 4–15% polyacrylamide gels (Bio-Rad Laboratories, Inc., Hercules, CA) and then transferred onto a nitrocellulose membrane (Bio-Rad Laboratories, Inc.). The membrane was blocked with SuperBlock™ Blocking Buffer (Thermo Fisher Scientific) for 1 hour at room temperature before being incubated overnight at 4°C with primary antibodies. Primary antibodies included ADRA2A (Thermo Fisher Scientific, Catalog #14266-1-AP, 1:500) and β-Actin (MilliporeSigma, A5441, 1:2000). Following primary antibody incubation, the membrane was treated with secondary ECL anti-rabbit or anti-mouse IgG HRP-linked antibodies (GE Healthcare, Pittsburgh, PA) for 1 hour at room temperature. Protein visualization was achieved using SuperSignal™ West Dura Extended Duration Substrate (Thermo Fisher Scientific).

### Statistical analysis

We conducted statistical analyses using GraphPad Prizm 9 (GraphPad Software, La Jolla, CA). Overall survival differences between patient groups were assessed using the Kaplan-Meier method, with significance determined by the log-rank test. Column differences were evaluated using the chi-square test. Differences between groups and their significance were evaluated using unpaired two-tailed Student’s t-tests (for two groups) or the ANOVA test (for three or more groups). Results are expressed as mean ± SD, with statistical significance defined as p < 0.05.

## Results

### Downregulated ADRA2A levels correlate with the basal-like/squamous subtype, advanced stage, and poor patient survival in PDAC

We aimed to identify potential suppressor genes implicated in the development of the basal-like/squamous subtype of PDAC (24). By conducting transcriptome analysis across three PDAC cohorts (Bailey cohort (3), Moffitt cohort (4), and NCI-UMD-German cohort (18)), we narrowed our candidate gene list down to 60 genes (24) (Figure 1A). Among these genes, *ADRA2A* held the highest ranks as the top 11 genes, displaying significantly lower hazard ratio (hazard ratio = 0.501; p < 0.05). Notably, ADRA2A has been recognized as a protein associated with cellular metabolism (27). *ADRA2A* exhibited a more substantial downregulation in tumor tissues compared to nontumor tissues. Building on these observations, we hypothesized that ADRA2A could inhibit PDAC progression, or the differentiation into the aggressive basal-like/squamous subtype, through metabolic reprogramming, prompting us to select it for in-depth investigation. We revealed that reduced *ADRA2A* transcript levels correlated with decreased patient survival. Additionally, *ADRA2A* transcripts displayed downregulation in tumor tissues compared to nontumor tissues in both our NCI-UMD-German cohort and the validation cohort (Moffitt cohort (4)) (Figure 1B-E). Furthermore, the downregulation of *ADRA2A* was associated with higher incidence of lymph node metastasis, advanced stages, and high-grade PDAC (Figure 1F). Immunohistochemistry (IHC) illustrated the presence of ADRA2A protein in the cytoplasm and the cellular membrane of acinar, endocrine, duct, and tumor cells, albeit at lower levels in tumor cells (Figure 2A-2C). Patients with ADRA2A-high PDAC associated with increased prognosis (Figure 2D). These findings support the hypothesis that upregulated ADRA2A may have a potential inhibitory role in PDAC whereas its diminished expression is associated with adverse patient survival and differentiation into the basal-like/squamous subtype.

**Figure 1.**
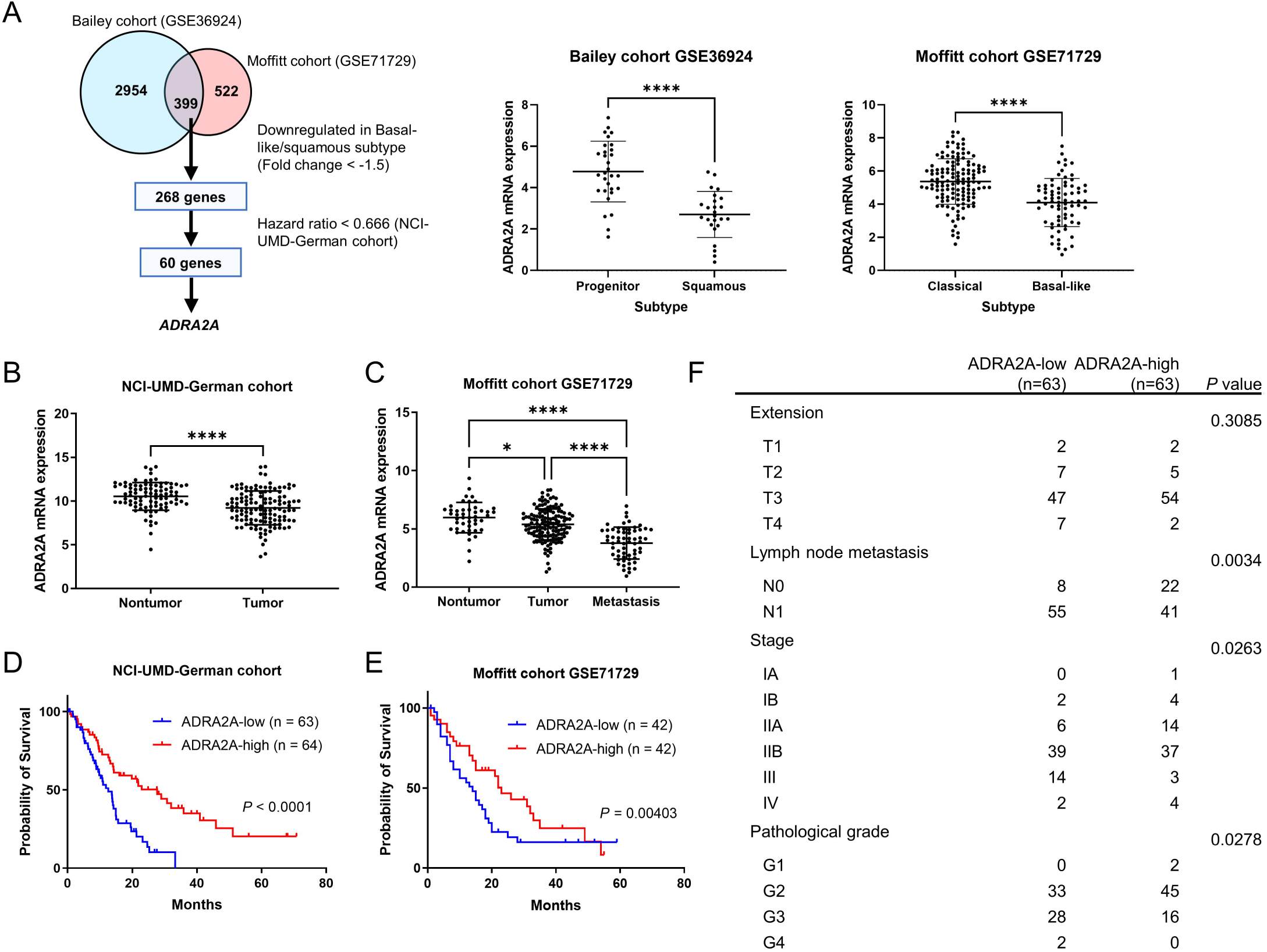
Transcriptome analysis identifies ADRA2A as a candidate suppressor of basal-like/squamous PDAC and a prognostic marker of patient survival. (A) Workflow for the identification of candidate suppressor genes in the basal-like/squamous subtype. Candidate genes were identified through the analysis of two cohorts (Bailey cohort (3) and Moffitt cohort (4)), followed by survival analysis in the NCI-UMD-German cohort (18). (B-E) Comparison of ADRA2A transcript levels in PDAC tumors and adjacent noncancerous tissues (B and C). ADRA2A expression is downregulated in tumors, as observed in both the NCI-UMD-German cohort (qPCR) and a validation cohort (Moffitt cohort (4)). Metastatic PDAC shows the lowest ADRA2A expression (C). Panels D and E show Kaplan-Meier plots and log-rank test results, depicting the association between decreased ADRA2A and decreased PDAC patient survival in the NCI-UMD-German and a validation cohort (Moffitt cohort (4)). The comparison was made between patients in the upper 33.3% tertile and the lower 33.3% tertile of ADRA2A expression in E. (F) Upregulation of ADRA2A is associated with negative lymph node metastasis, earlier-stage, and low-grade PDAC (qPCR). The grades are categorized as follows: Grade 1 (well differentiated), Grade 2 (moderately differentiated), Grade 3 (poorly differentiated), and Grade 4 (undifferentiated). Data represent mean ± SD. *p ≤ 0.05, **p ≤ 0.01, ***p ≤ 0.001, ****p ≤ 0.0001 by unpaired two-tailed Student’s t-test or one-way ANOVA. Chi-square test for F.

**Figure 2.**
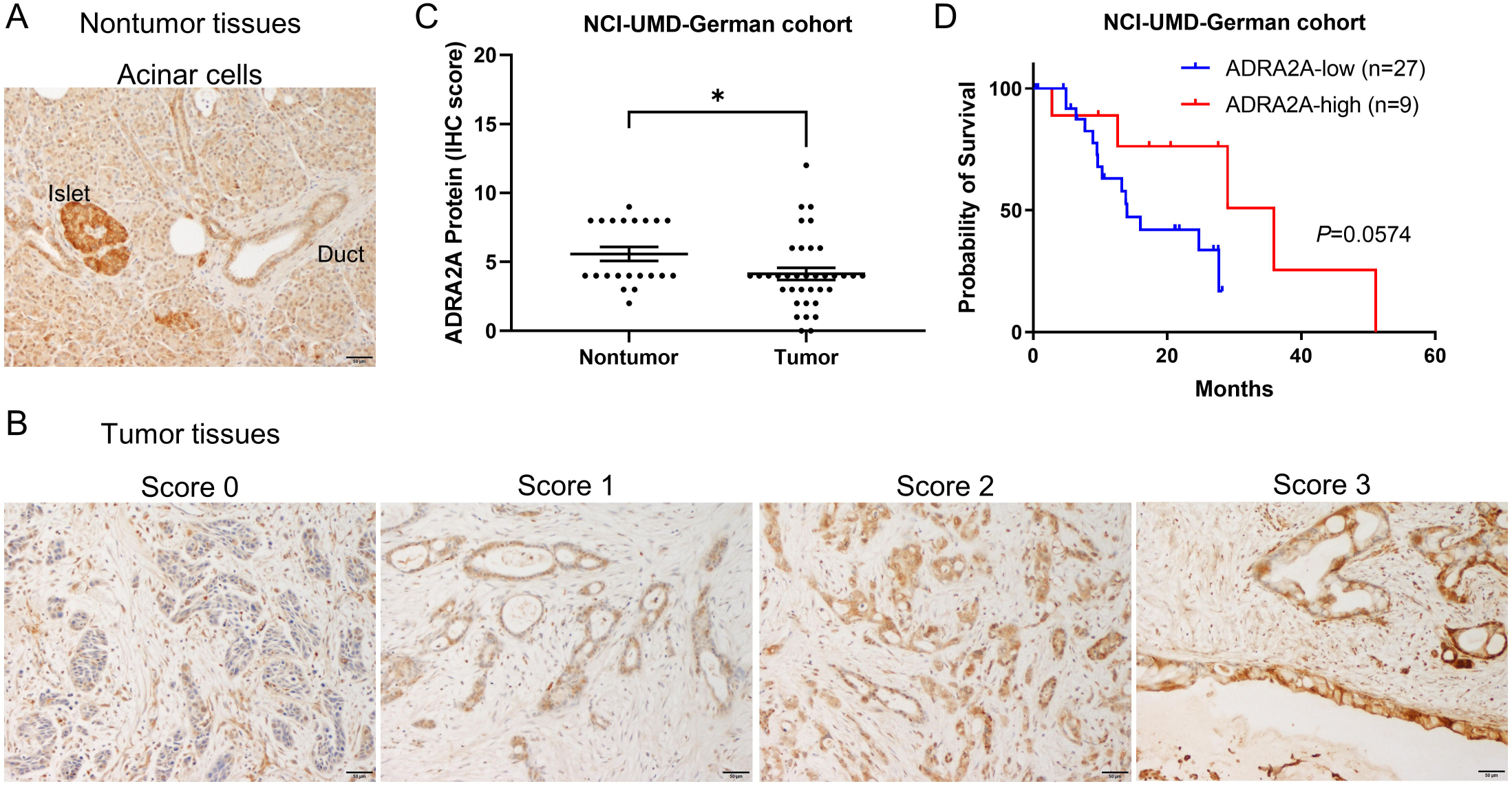
Downregulation of ADRA2A protein in PDAC tumors is correlated with patient poor prognosis in the NCI-UMD-German cohort. (A and B) IHC of ADRA2A in tumor and nontumor tissue sections of PDAC patients. ADRA2A protein is detected in the cytoplasm and the cellular membrane, as shown by the brown DAB-based IHC in the acinar cells, the endocrine cells in pancreatic islets, and the tumor cells. (A) is a representative ADRA2A protein staining in noncancerous acinar cells (intensity score 2 in acinar cells and score 3 in islets). (B) is ADRA2A protein in representative tumor sections (scores 0-3). The staining strength is categorized as follows: score 0 (unstained), score 1 (weak), score 2 (moderate), and score 3 (strong). (C) Downregulation of ADRA2A in tumor cells when compared to the nontumor acinar cells. (D) The result shows the tendency of ADRA2A protein downregulation in PDAC tumors correlated with patient poor prognosis in the NCI-UMD-German cohort. Scale bar is 50 μm. More details can be found in Materials and Methods. Data represent mean ± SD. *p ≤ 0.05 by unpaired two-tailed Student’s t-test. The Kaplan-Meier method was employed to assess the difference in overall survival between patient groups, and significance was determined using the log-rank test. DAB; 3, 3’-diaminobenzidine, IHC; immunohistochemistry.

### Upregulation of ADRA2A upregulates the classical/progenitor subtype differentiation in PDAC

We next investigated whether ADRA2A expression would promote the characteristics associated with the classical/progenitor subtype, thereby reducing disease aggressiveness. In our previous study, we categorized 175 patients from our NCI-UMD-German cohort into molecular subtypes: unclassified (n = 32), classical/progenitor (n = 95), and basal-like/squamous (n = 48) (17). We used a gene set from Moffitt et al. (4) to define these subtypes, where the classical/progenitor and the basal-like/squamous subtypes upregulated specific genes, while the unclassified subtype downregulated these specific genes (17). We showed that pathways associated with cellular movement were activated in the basal-like/squamous subtype (17). Extending this work, we found that tumors categorized as the basal-like/squamous subtype showed lower levels of *ADRA2A* mRNA expression compared to the other subtypes (Figures S1A). Gene set enrichment analysis (GSEA) revealed that *ADRA2A*-high PDAC tumors in the NCI-UMD-German cohort downregulated the pathways associated with the basal-like/squamous subtype, including hypoxia, epithelial-mesenchymal transition (EMT), and MYC signaling pathways (3,17,28) (Figure 3A and 3B). Simultaneously, *ADRA2A*-high tumors downregulated the basal-like gene signature (gene set upregulated in basal-like tumors (4,29)) (Figure 3B and Figure S1B), while upregulated the classical gene expression signature (4) (Figure 3B). Pathway enrichment analysis using the Ingenuity Pathways Analysis (IPA) revealed that *ADRA2A*-high PDAC tumors downregulated pathways related to cellular movement (Figure 3C). To validate these observations, we assessed ADRA2A expression in human PDAC cell lines, revealing that ADRA2A expression remained consistently low to undetectable across all the PDAC cell lines (Figure S1C and S1D). We selected CFPAC-1 and Panc 10.05 cells to establish cell lines with *ADRA2A* transgene expression. The upregulated ADRA2A expression was confirmed at both the mRNA and protein levels in these cell lines (Figure S1E-S1H). GSEA and IPA revealed concordance among enriched pathways in PDAC cells with *ADRA2A* transgene expression and *ADRA2A*-high PDAC tumors in the NCI-UMD-German cohort, highlighting the downregulation of pathways associated with the basal-like/squamous subtype and the pathways linked to cellular movement (Figure 3D-3F). In contrast, PDAC cells with *ADRA2A* transgene expression upregulated the classical gene signature (Figure 3E). IPA’s upstream analysis revealed that ADRA2A-high conditions inhibited various regulators, including TGF-β, HIF-1A, protein kinase C (PKC), and the mitogen-activated protein kinase (MAPK) pathways (Table S1). These findings align with earlier observations regarding the basal-like/squamous subtype (3) and ADRA2A (30–33). To further investigate ADRA2A’s role, we conducted additional *in vitro* experiments using these ADRA2A-overexpressing PDAC cells. ADRA2A expression significantly inhibited the invasion of PDAC cells (Figure 3G and 3H). Additionally, an ADRA2A agonist, clonidine, inhibited the PDAC cell invasion (Figure 3I and 3J). Collectively, these findings suggest that ADRA2A plays a role in driving the classical/progenitor subtype differentiation in PDAC.

**Figure 3.**
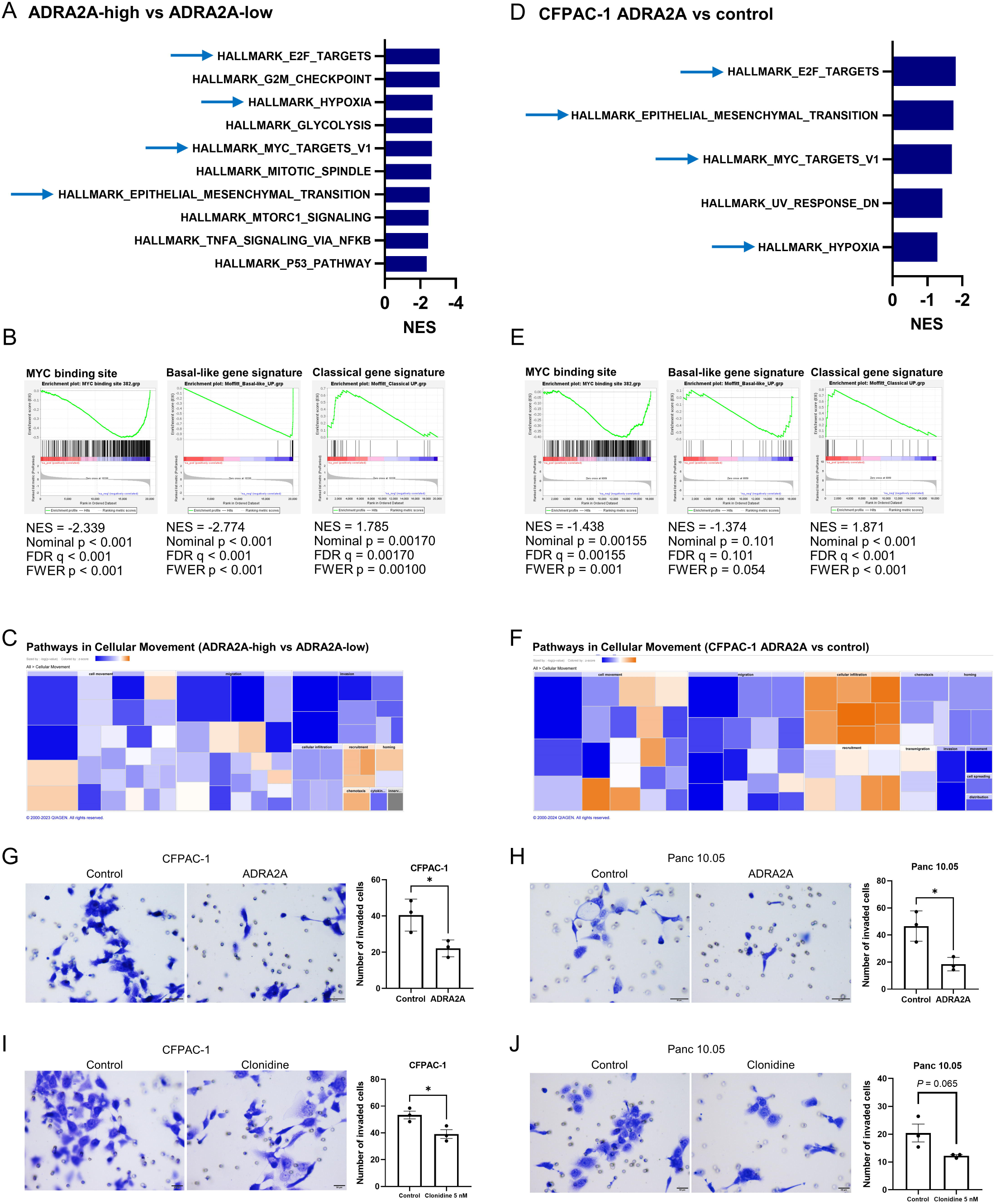
ADRA2A suppresses the basal-like/squamous subtype differentiation in PDAC. (A-F) Transcriptome analyses of patient PDAC and Panc 10.05 human PDAC cells using Gene Set Enrichment Analysis (GSEA). The comparison involved *ADRA2A*-high versus *ADRA2A*-low PDAC tumors and CFPAC-1 cells expressing the ADRA2A transgene (ADRA2A-high) versus vector control (ADRA2A-low). GSEA reveals the association of ADRA2A-high conditions with the downregulated pathways linked to the basal-like/squamous subtype in both patient PDAC and cultured cells, particularly hypoxia, MYC, and epithelial-mesenchymal transition (A and D). ADRA2A-high conditions downregulated MYC signaling (MYC binding site (17,28)) and the basal-like gene expression signature (4), while upregulated the classical gene expression signature (4) (B and E). (C and F) Ingenuity Pathway Analysis (IPA) shows the Z score-based enrichment heatmap (orange color; upregulation, blue color; downregulation) of pathways related to cellular movement, showcasing the common downregulation of these pathways in the ADRA2A-high conditions. (G and H) PDAC cells overexpressing ADRA2A display diminished invasion capabilities compared to vector control cells. (I and J) PDAC cells diminished invasion capabilities with supplementation of ADRA2A agonist, clonidine. Data represent mean ± SD of three replicates. *p ≤ 0.05 by unpaired two-tailed Student’s t-test.

### Downregulation of amino acid and carnitine/acylcarnitine metabolism in ADRA2A-high PDAC

Expanding our investigation to include metabolome analysis, we employed transcriptome and metabolome analyses on a cohort of 50 patients from our NCI-UMD-German cohort (23). We identified 7 significantly upregulated and 76 downregulated metabolites in *ADRA2A*-high tumors when compared to *ADRA2A*-low tumors (Figure 4A). Pathway enrichment analysis using MetaboAnalyst 5.0 (25) with these differential metabolites as input revealed the downregulation of amino acid metabolism in *ADRA2A*-high tumors (Figure 4B). To delve deeper into the roles of these metabolites, we conducted a comprehensive metabolome analysis in Panc 10.05 cells. We found that *ADRA2A* transgene overexpression resulted in the upregulation of 30 metabolites and the downregulation of 265 metabolites when compared to vector control cells (Figure 4C). Applying pathway enrichment analysis using MetaboAnalyst 5.0 (25), we noticed that ADRA2A-overexpressing Panc 10.05 cells exhibited diminished amino acid metabolism (Figure 4D). Our findings corroborate earlier research conducted by our group, indicating that the basal-like/squamous subtype fosters amino acid metabolism, whereas the classical/progenitor subtype suppresses it (17,23,24). Additionally, *ADRA2A*-high tumors and ADRA2A-overexpressing Panc 10.05 cells showed the downregulation of carnitine/acylcarnitines. Specifically, one acylcarnitine was upregulated, and 22 acylcarnitines were downregulated in *ADRA2A*-high tumors, while one acylcarnitine was upregulated, and 25 carnitine/acylcarnitines were downregulated in ADRA2A-overexpressing Panc 10.05 cells (Table S2). Our previous study unveiled that these metabolites were upregulated in the basal-like/squamous subtype and correlated with decreased patient survival (17), supporting the current findings. As illustrated in Figure 3, ADRA2A inhibits MYC signaling. To further explore the metabolic reprogramming linked to ADRA2A’s downstream, we investigated amino acid metabolism in PDAC cells supplemented with a MYC inhibitor. Our results indicated that MYC inhibition downregulated amino acid metabolism (Figure 5, Figure S2, and Table S3). These results suggest that ADRA2A contributes to the metabolic characteristics of the classical/progenitor subtype of PDAC.

**Figure 4.**
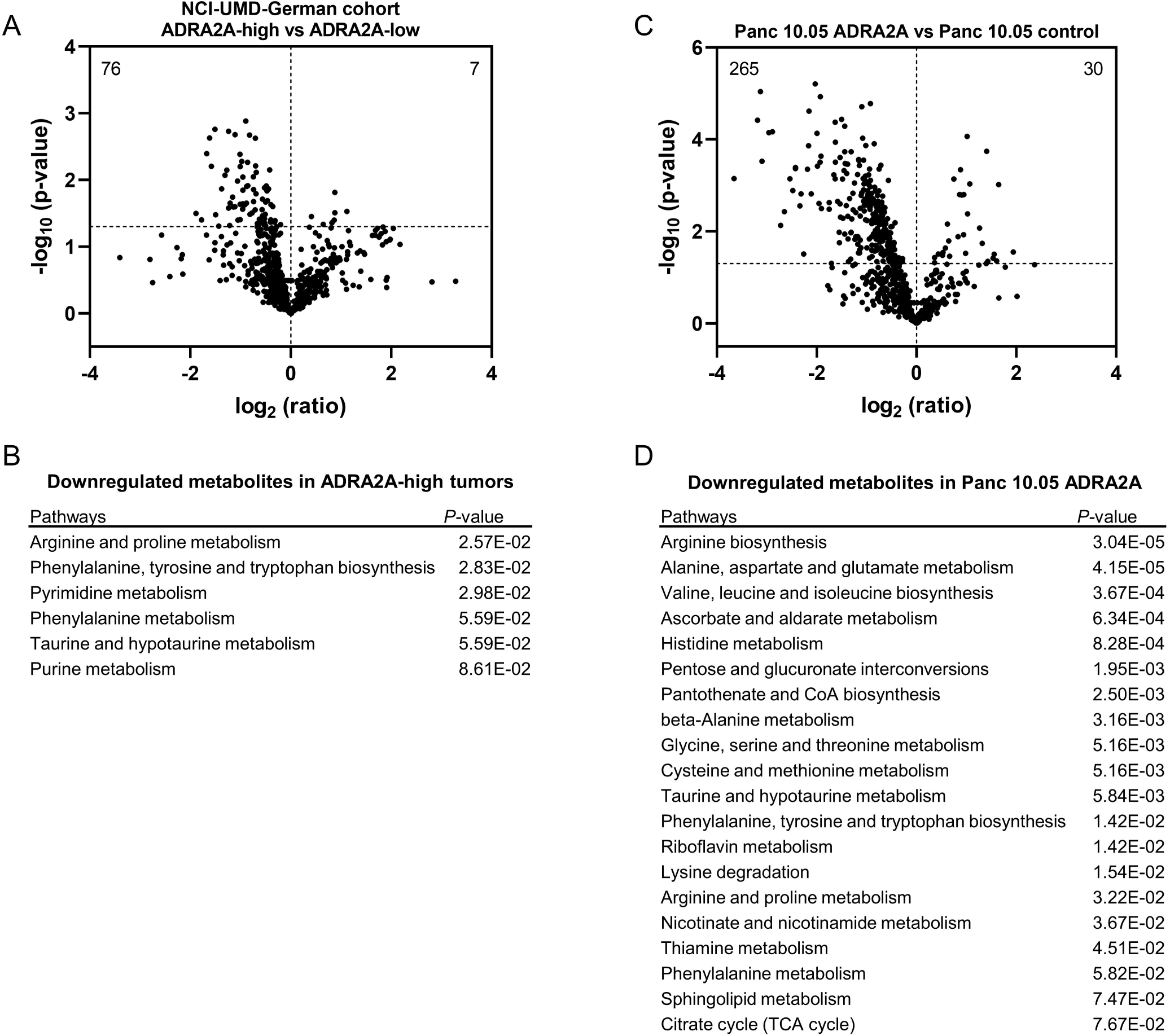
Downregulation of amino acid and carnitine/acylcarnitine metabolism in ADRA2A-high PDAC. Metabolome analysis conducted on PDAC tumors (NCI-UMD-German cohort) and PDAC cells. The results depict comparisons between *ADRA2A*-high (n = 25) and *ADRA2A*-low (n = 25) PDAC tumors, and Panc 10.05 with ADRA2A transgene expression (ADRA2A-high) and Panc 10.05 with empty control vector (ADRA2A-low). (A) Volcano plot displaying metabolites with differential abundance in *ADRA2A*-high versus *ADRA2A*-low tumors. 7 metabolites are significantly upregulated and 76 are downregulated in *ADRA2A*-high tumors (p < 0.05). The dotted line indicates –log_10_ (p = 0.05). (B) Pathway enrichment analysis with differential metabolites using MetaboAnalyst 5.0 (25), highlighting downregulated amino acid metabolism in *ADRA2A*-high tumors. (C) Volcano plot illustrating metabolites with differential abundance in ADRA2A-overexpressing versus vector control Panc 10.05 cells. 30 metabolites are significantly upregulated and 265 are downregulated in ADRA2A-overexpressing Panc 10.05 cells. The dotted line indicates –log_10_ (p = 0.05). (D) Pathway enrichment analysis with differential metabolites using MetaboAnalyst 5.0 (25), demonstrating downregulated amino acid metabolism in ADRA2A-overexpressing Panc 10.05 cells.

**Figure 5.**
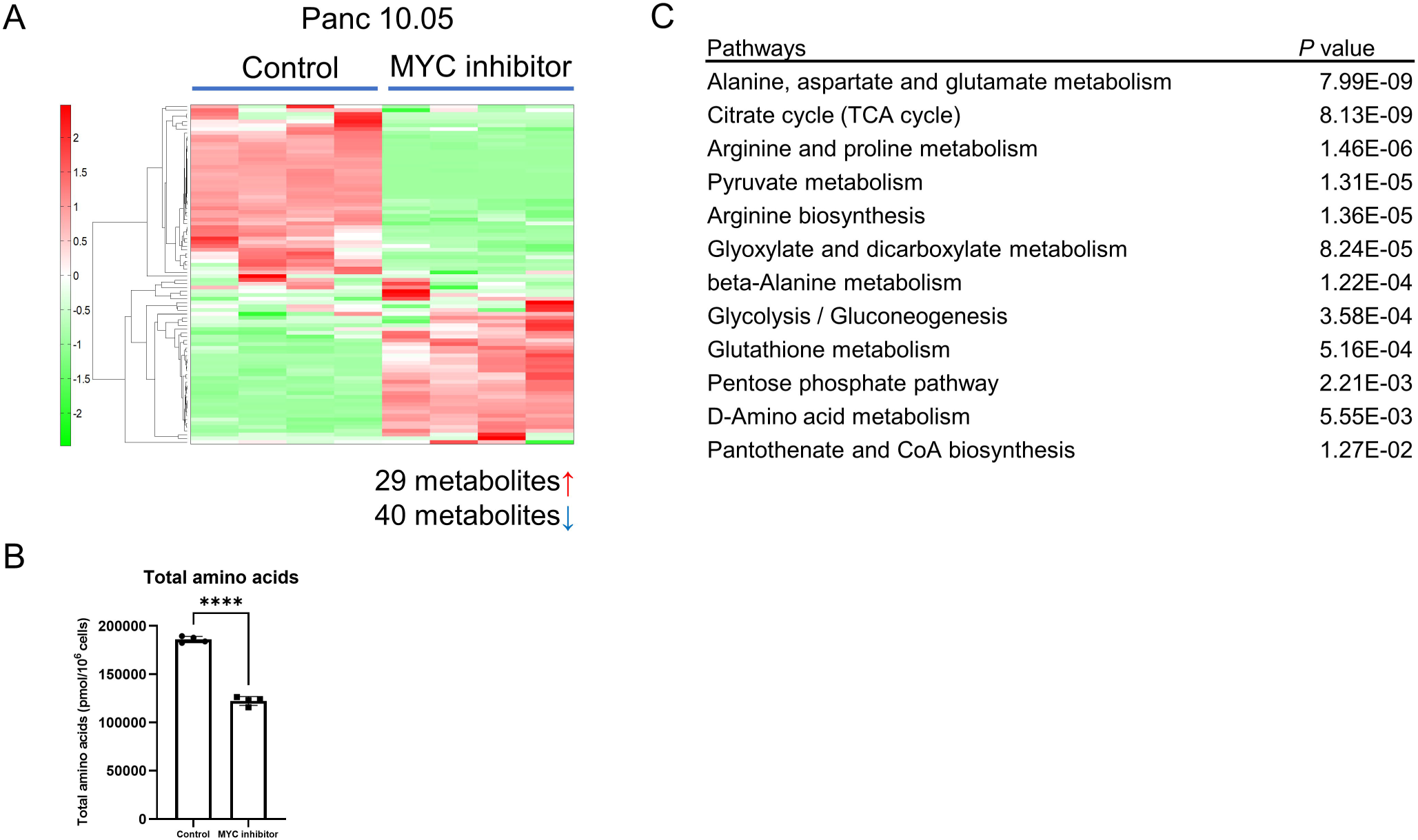
Downregulation of amino acid metabolism in Panc 10.05 cells with a MYC inhibitor. (A and B) Metabolome analysis covering 116 cancer-related metabolites in Panc 10.05 cells with a MYC inhibitor (10058-F4, 100 μM). In Panc 10.05 cells with a MYC inhibitor, 40 metabolites were significantly downregulated when compared with solvent control cells (p < 0.05). The graphs illustrate the decrease in amino acid metabolism. Data are presented as mean ± SD. (C) Pathway enrichment scores using MetaboAnalyst 5.0 (25) with the significantly decreased 40 input metabolites, indicating that amino acid metabolism were downregulated in Panc 10.05 cells with a MYC inhibitor. Data represent mean ± SD of four replicates. ****p ≤ 0.0001 by unpaired two-tailed Student’s t-test.

## Discussion

In this study, we observed that the upregulation of ADRA2A is associated with the differentiation into the classical/progenitor subtype and predicts a favorable prognosis for patients with PDAC. Elevated ADRA2A induced changes in cancer cell metabolism, resulting in reduced invasion in PDAC cells. The upregulation of ADRA2A led to the downregulation of MYC signaling and the increase of the classical/progenitor gene signature in patient PDAC samples and PDAC cells.

Adrenoceptor alpha 2A (ADRA2A) is known for its involvement in the regulation of neurotransmitter release and modulation of sympathetic nervous system activity (30,31). Its functions extend beyond the central nervous system, encompassing diverse physiological functions such as cardiovascular regulation and smooth muscle contraction (30,31). For cellular responses, ADRA2A inhibits adenylate cyclase, leading to decreased cyclic adenosine monophosphate (cAMP) levels and modulation of various intracellular pathways, including protein kinase A (PKA), PKC, and the MAPK pathways (30–33). ADRA2A also plays a role in metabolism, exerting control over lipid and glucose homeostasis in adipose tissue and the liver (27,34,35). In cancers, ADRA2A exhibits a complex interplay, with reports indicating both tumor-promoting and tumor-suppressing effects across various cancer types. In breast cancer, while patients with ADRA2A-high breast cancer showed a favorable prognosis (36), it may promote metastasis (37). ADRA2A promotes senescence and apoptosis through the inhibition of the PI3K/Akt/mTOR pathway in cervical cancer (38). Activation of ADRA2A promotes the chemosensitivity of carboplatin in ovarian cancer cells (32). A recent study has shown that ADRA2A agonist highlights cancer immunotherapy, suppressing tumor growth in colorectal cancer and melanoma (39). ADRA2A’s impact on critical cancer processes underscores its significance as a potential therapeutic target. Understanding the diverse functions of ADRA2A is crucial for elucidating its therapeutic potential in metabolic disorders and cancer treatment. While previous studies have highlighted ADRA2A’s involvement in cancers, its significance in PDAC remains relatively understudied. Our study bridges this gap by uncovering the association between ADRA2A downregulation and the basal-like/squamous subtype of PDAC, shedding light on its potential role as a prognostic marker and therapeutic target in this aggressive malignancy.

In a previous study, we clustered 175 patients in our NCI-UMD-German cohort into unclassified, classical/progenitor, and basal-like/squamous subtypes (17) using a gene set developed by Moffitt et al. (4). This gene set comprises the genes upregulated in the classical subtype and those upregulated in the basal-like subtype for molecular subtype classification. The classical/progenitor and basal-like/squamous subtypes upregulated specific genes, while the unclassified subtype downregulated these genes (17). The basal-like/squamous subtype upregulated amino acid and carnitine/acylcarnitine metabolism compared to the classical/progenitor subtype, while unclassified subtype upregulated lipogenesis. Recognizing the biological similarity, we grouped unclassified and classical/progenitor subtypes into a unified group termed “unified classical/progenitor subtype” (23). LIM Domain Only 3 (LMO3) downregulated both basal-like genes and classical genes developed by Moffitt et al. (4) to induce the unified classical/progenitor subtype (23). However, the current study revealed that ADRA2A upregulates the classical genes while downregulating the basal-like genes to promote classical/progenitor subtype differentiation. Furthermore, ADRA2A downregulated amino acid and carnitine/acylcarnitine metabolism. Thus, ADRA2A plays a role inducing the pure classical/progenitor subtype defined in our first study (17), without inducing the unclassified subtype. This difference between ADRA2A and LMO3 may contribute to marker and metabolic explorations for PDAC molecular subtypes.

Cancer cells exhibit metabolic reprogramming characterized by increased glycolysis, lipid synthesis, and amino acid production (8,9). Adrenoceptor beta 2 (ADRB2) is a member of adrenergic receptor family, sharing similar functions with ADRA2A. A recent study revealed that ADRB2 agonists downregulated the MYC pathway and HIF-1A, inducing specific metabolic reprogramming in PDAC (40). These reprogramming include the upregulation of glycolytic metabolism and changes in acylcarnitine levels (40), ultimately leading to the suppression of PDAC progression. These findings parallel our observations regarding ADRA2A in this study.

Our previous study unveiled that MYC induces the basal-like/squamous gene signature (17). In the current study, we observed that ADRA2A downregulated MYC signaling and the MAPK and PKC pathways in the transcriptome analysis. The suppression of the basal-like/squamous subtype differentiation by ADRA2A appeared to stem from the downregulation of MYC. This downregulation of MYC may be attributed to the reduction in MAPK and PKC pathway activity. However, our study is constrained by the absence of mechanistic analyses exploring the downstream pathways of ADRA2A, including the MAPK and PKC pathways (30–33). While our research demonstrated upregulation of ADRA2A resulted in reduced amino acid metabolism, reduced cellular activity, downregulation of MYC signaling, induction of the classical/progenitor subtype, and extended patient survival in PDAC, our plan is to address the limitation through comprehensive analysis in future studies. Further exploration holds the potential to provide valuable insights for targeting ADRA2A-downstream pathways. Specifically, we aim to elucidate how ADRA2A downstream pathways regulate metabolic reprogramming in PDAC. Additionally, this study focused on gain-of-function experiments because all the PDAC cell lines we utilized exhibited downregulated ADRA2A expression. Future investigations should include mechanistic studies and loss-of-function experiments to provide a more comprehensive understanding of ADRA2A’s role in PDAC.

In summary, our findings indicate that ADRA2A expression suppresses transcriptome features of the basal-like/squamous PDAC subtype, while promoting those of the classical/progenitor subtype, leading to decreased PDAC aggressiveness. Furthermore, ADRA2A promotes a distinct metabolic profile characterized by downregulated amino acid and carnitine/acylcarnitine metabolism, a hallmark of the classical/progenitor PDAC subtype.

## Abbreviations

ADEX; aberrantly differentiated endocrine exocrine, ADRA2A; adrenoceptor alpha 2A, DAB; 3, 3’-diaminobenzidine, GSEA; Gene Set Enrichment Analysis, IHC; immunohistochemistry, IPA; Ingenuity pathway analysis, LMO3; LIM Domain Only 3, PDAC; pancreatic ductal adenocarcinoma, STR; short tandem repeat

## Acknowledgments

The authors would like to extend their gratitude to the dedicated staff and study coordinators at the University of Maryland School of Medicine for their invaluable assistance in obtaining clinical biospecimens and patient data. Additionally, we extend our appreciation to the Department of General, Visceral, and Pediatric Surgery at the University Medical Center Göttingen, Göttingen, Germany, for their generous support in handling clinical samples. Special thanks are extended to the dedicated team at the NCI-CCR Sequencing Facility Frederick for their invaluable contributions to this research. We extend our heartfelt appreciation to the late S. Perwez Hussain, the last author of this manuscript, for his contributions to conceptualization, writing (review & editing), supervision, and funding acquisition for this study. In recognition of his significant efforts and steadfast dedication to this project, we respectfully include S. Perwez Hussain as a co-author of this paper. His enduring legacy continues to serve as an inspiration to our endeavors.

## Data availability

The RNA sequencing data were deposited in the NCBI’s Gene Expression Omnibus (GEO) database under accession number GSE259390: ADRA2A is a suppressor of the basal-like/squamous PDAC subtype and reduces disease aggressiveness of pancreatic cancer. To review GEO accession GSE259390. The transcriptome and metabolome data used in this study have been deposited in the Open Science Framework (https://osf.io). Other data that support the findings of this study are available from the corresponding author upon request.

## Author Contributions

**Paloma Moreno:** Methodology, Resources, Investigation, Project Administration, Writing – Review & Editing. **Yuuki Ohara:** Conceptualization, Methodology, Resources, Investigation, Writing – Original Draft, Writing – Review & Editing, Project Administration. **Amanda J. Craig**: Methodology, Data Curation, Writing – Review & Editing, Project Administration. **Huaitian Liu:** Data Curation, Writing – Review & Editing, Project Administration. **Shouhui Yang:** Methodology, Writing – Review & Editing, Project Administration. **Lin Zhang**: Project Administration. **Gatikrushna Panigrahi:** Project Administration. **Tiffany H. Dorsey:** Project Administration. **Helen Cawley:** Project Administration. **Azadeh Azizian:** Resources, Project Administration. **Jochen Gaedcke:** Resources, Project Administration. **Michael Ghadimi:** Resources, Project Administration. **Nader Hanna:** Resources, Project Administration. **S. Perwez Hussain:** Conceptualization, Writing – Review & Editing, Supervision, Funding Acquisition. The work reported in the paper has been performed by the authors, unless clearly specified in the text.

## Supplementary figure legends

**Supplementary Figure 1.**
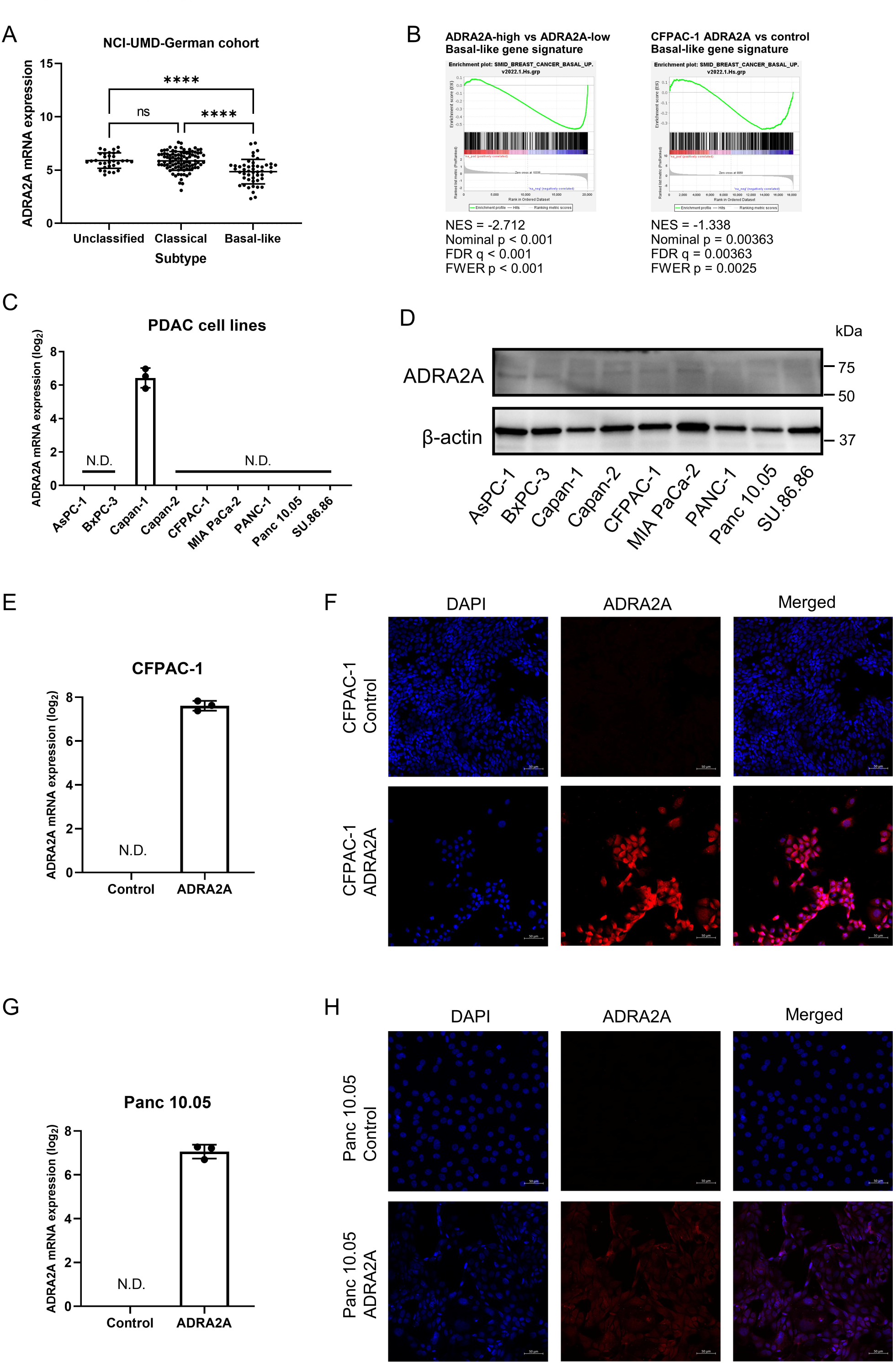
ADRA2A expression and analysis by molecular subtypes in PDAC. (A) *ADRA2A* mRNA expression by molecular subtypes in the NCI-UMD-German cohort. (B) Gene Set Enrichment Analysis (GSEA) suggests that elevated ADRA2A expression leads to the downregulation of the predefined basal-like gene expression profile (29). (C and D) Endogenous levels of ADRA2A mRNA and protein in various human PDAC cell lines, revealing consistent suppression of ADRA2A expression across all examined PDAC cell lines. (E-H) Confirmation of *ADRA2A* transgene overexpression at the mRNA (qPCR) and protein levels in CFPAC-1 and Panc 10.05 cells. Scale bar is 50 μm. Data represent mean ± SD with ANOVA. ****p ≤ 0.0001. N.D.; not detected.

**Supplementary Figure 2.**
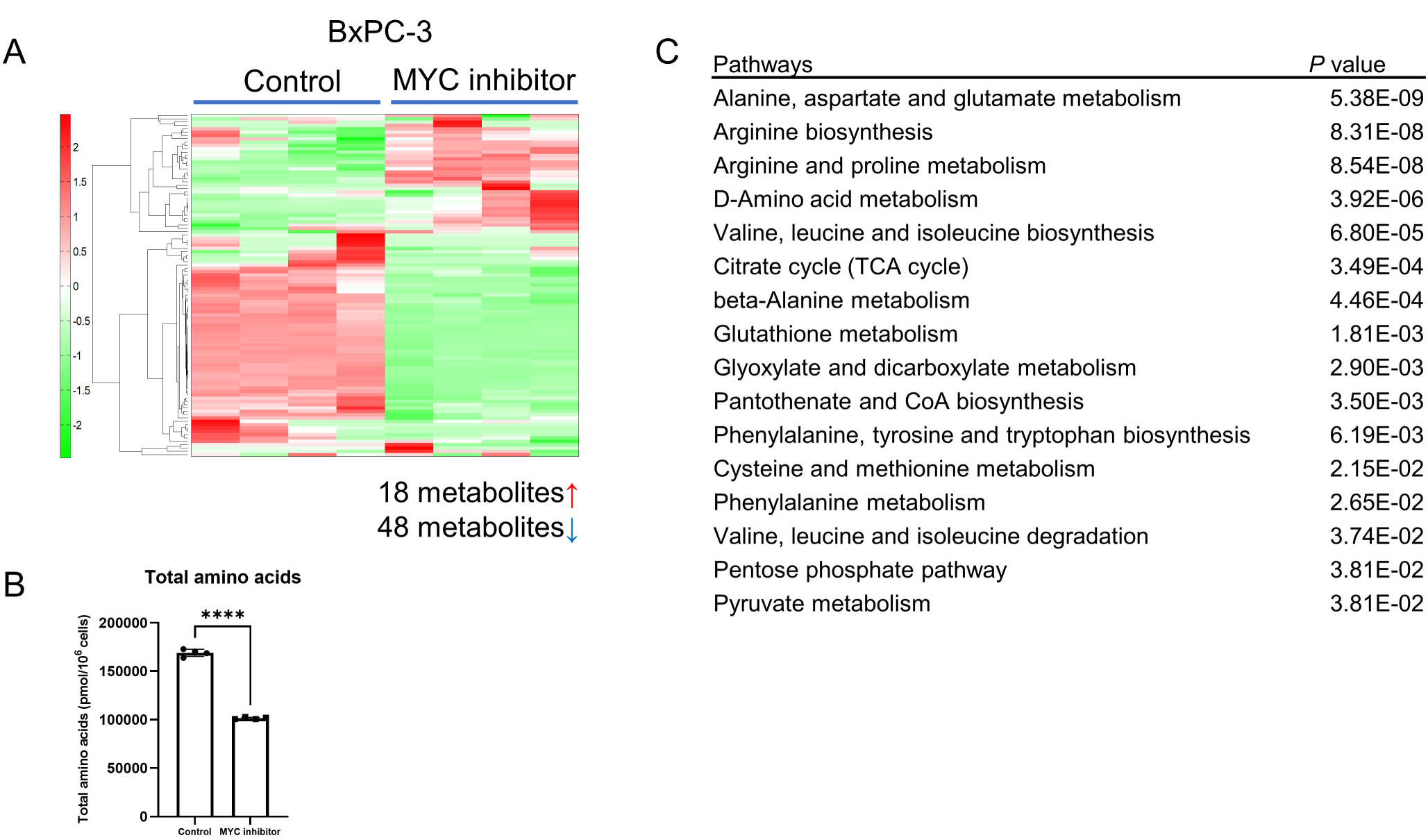
Downregulation of amino acid metabolism in PDAC cells with a MYC inhibitor. (A and B) Metabolome analysis covering 116 cancer-related metabolites in BxPC-3 cells with a MYC inhibitor (10058-F4, 100 μM). In BxPC-3 cells with a MYC inhibitor, 48 metabolites were significantly downregulated when compared with solvent control cells (p < 0.05). The graphs illustrate the decrease in amino acid metabolism. Data are presented as mean ± SD. (C) Pathway enrichment scores using MetaboAnalyst 5.0 (25) with the significantly decreased 48 input metabolites, indicating that amino acid metabolism were downregulated in BxPC-3 cells with a MYC inhibitor. Data represent mean ± SD of four replicates. ****p ≤ 0.0001 by unpaired two-tailed Student’s t-test.

